# A probiotic bacterium modulates antitumor γδ T-cell responses in lung cancer

**DOI:** 10.1101/2025.07.16.664923

**Authors:** Yoshihiko Goto, Garry Dolton, Hannah Thomas, Théo Morin, Yuka Tajima, Kosuke Imamura, Shinya Sakata, Kentaro Oka, Atsushi Hayashi, Motomichi Takahashi, Takamasa Ueno, Takuro Sakagami, Yusuke Tomita, Andrew K. Sewell, Chihiro Motozono

## Abstract

The link between the intestinal microflora and cancer outcomes has been recognized for over a decade. Several recent studies have demonstrated that the gut microbiome is associated with the efficiency of T-cell checkpoint blockade therapy for cancer raising interest in strategies to harness this effect via consumption of live microorganisms (probiotics). The probiotic *Clostridium butyricum* strain MIYAIRI 588 (CBM588) enhances both response rates and overall survival in patients receiving immune checkpoint blockade therapy for non-small cell lung cancer and metastatic renal cell carcinoma but the mechanism underpinning this benefit remains unclear. Here, we show that CBM588 spores induce a population of Vγ9Vδ2 T-cells from the peripheral blood of healthy donors and lung cancer patients. A subset of these T-cells responded to, and directly lysed, cancer cell lines via a butyrophilin 3A-dependant mechanism. In patients taking CBM588 alongside checkpoint blockade, peripheral Vδ2⁺ T-cells expressed the activation marker CD69 more frequently than those receiving checkpoint blockade alone and the frequency of Vδ2⁺CD69⁺ cells increased following initiation of CBM588 treatment (p = 0.0041). Our findings provide a potential mechanism by which the manipulation of the intestinal microflora might directly contribute to cancer prognosis by activating immune effector cells with intrinsic anticancer properties.

## INTRODUCTION

Many cancers are associated with an altered composition of commensal microbiota, raising debate over whether this dysbiosis is a cause or effect of malignancy. Evidence for a causal role has emerged from gnotobiotic mouse models, which show that the microbiota can profoundly influence cancer susceptibility and progression (1). Early animal studies demonstrated that the microbiome modulates the anticancer efficacy of cytotoxic agents such as cyclophosphamide and oxaliplatin (2, 3). More recently, multiple studies have shown that the gut microbiome also impacts the effectiveness of T-cell checkpoint blockade therapy across several cancer types (4–7). The strong association between the gut microbiota and cancer treatment outcomes has resulted in fervent interest in harnessing this affect through the use of probiotics (8). A notable example is *Clostridium butyricum* MIYAIRI 588 (CBM588), an anaerobic probiotic. In retrospective studies at Kumamoto University Hospital, CBM588 was administered alongside checkpoint blockade therapy for non-small cell lung cancer (NSCLC), resulting in significantly improved progression-free survival (PFS; 250 vs. 101 days, *p* = 0.009) and overall survival (OS; median not reached vs. 361 days, *p* = 0.005) compared to patients not receiving the probiotic (9, 10). Additional support comes from a randomized phase 1 trial at City of Hope Medical Centre (NCT03829111), in which daily CBM588 improved response rates and extended PFS in patients with metastatic renal cell carcinoma (mRCC) treated with nivolumab and ipilimumab from 2.5 months to over one year (*p* = 0.001) (11). A second prospective study in mRCC (NCT05122546) reported significantly enhanced objective response rates (*p* = 0.01) and improved six-month PFS in patients receiving CBM588 alongside cabozantinib and nivolumab (12), generating further enthusiasm for this approach.

These clinical benefits have prompted efforts to validate the findings in larger patient cohorts and to understand the mechanisms by which probiotics such as CBM588 enhance cancer outcomes. We hypothesized that CBM588 might activate unconventional anticancer T-cells. Here, we show that CBM588 probiotic tablets containing *Clostridium butyricum* spores and purified spores alone induced a population of Vγ9Vδ2 T-cells from peripheral blood mononuclear cells (PBMCs) of healthy donors from the UK and Japan. A subset of these Vδ2⁺ T-cells responded to a lung cancer cell line via a butyrophilin 3A (BTN3A)-dependent mechanism and were capable of lysing multiple cancer cell lines *in vitro*. Notably, in a retrospective clinical cohort, the activation status of peripheral Vγ9Vδ2 T-cells measured by CD69 expression was higher in lung cancer patients taking CBM588 than in those receiving checkpoint blockade alone, and the frequency of Vδ2⁺CD69⁺ cells increased following initiation of CBM588. These results suggest a mechanism by which microbial probiotics can directly stimulate anticancer T-cell responses and potentially improve cancer prognosis.

## RESULTS

### The probiotic CBM588 induces Vγ9Vδ2 T-cells from healthy PBMC *in vitro*

We hypothesized that CBM588 may exert its striking clinical benefit for NSCLC (9, 10) by inducing T-cells capable of recognizing cancer cells. To test this, we stimulated PBMCs from healthy donors with a suspension of CBM588 tablets for 7 and 12 days, as described in the Materials and Methods. Flow cytometry analysis (gating strategy: **Supplementary Fig. 1A**) revealed that CBM588 induced a dose-dependent increase in γδ TCR⁺ T-cells, accompanied by a relative decrease in αβ TCR⁺ T-cells within the CD3⁺ population (**Fig. 1A**). The frequency of γδ TCR⁺ T-cells closely matched that of Vδ2⁺ T-cells, suggesting that CBM588 preferentially expanded the Vδ2⁺ subset without broadly increasing other γδ T-cell populations (**Fig. 1A**). Further staining confirmed that the Vδ2⁺ T-cells co-expressed Vγ9, identifying them as Vγ9Vδ2 T-cells (**Supplementary Fig. 2B**). Analysis of αβ TCR⁺ T-cells showed a slight shift in co-receptor expression, with a subtle decrease in CD4⁺ and increase in CD8⁺ T-cells, while the proportions of double-positive and double-negative subsets remained unchanged (**Supplementary Fig. 2C**). Among the CBM588-induced Vδ2⁺ T-cells, the majority were CD4^neg^/CD8^neg^, with approximately 15% expressing CD8 (**Supplementary Fig. 2D**).

**Figure 1:**
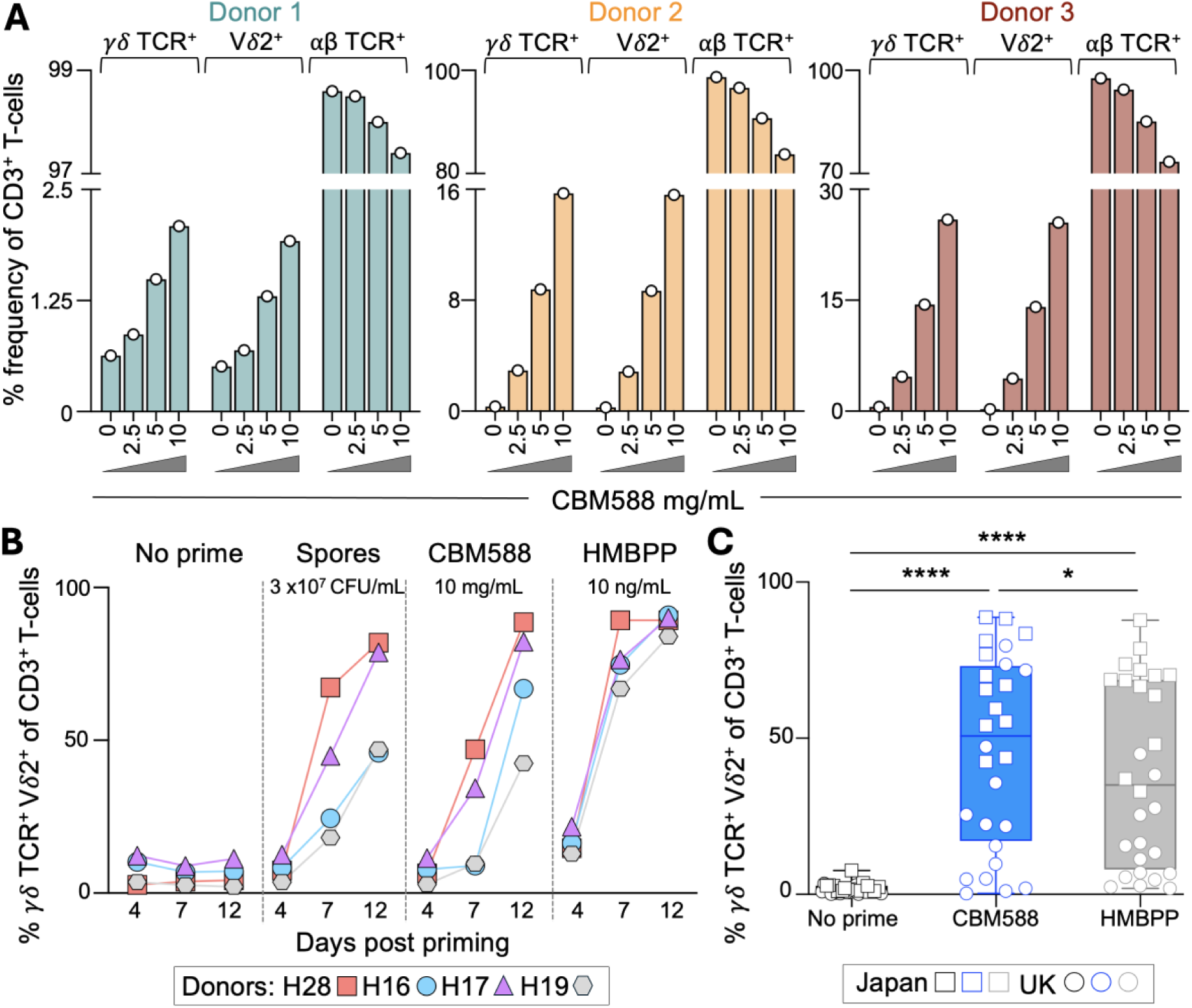
In vitro induction of γδ TCR^+^ T-cells by CBM588 tablet or bacterium spores in healthy donors. **A**. PBMCs from healthy donors were stimulated for 12 days with a suspension of CBM588 at the concentrations shown. Flow cytometry gating strategies, αβ TCR^+^ T-cell subset definitions, and co-receptor and Vγ9 expression within the Vδ2^+^ T-cell population are shown in **Supplementary Fig. B.** Frequency of γδ TCR^+^ Vδ2^+^ T-cells among CD3^+^ T-cells following stimulation of PBMCs from four healthy donors with CBM588 spores (3×10⁷ CFU/mL), CBM588 tablet suspension (10 mg/mL), or HMBPP (10 ng/mL) at days 4, 7 and 12 post stimulation. Results at other concentrations of spore, CBM588 tablet and HMBPP are shown in **Supplementary Fig. 2**, alongside αβ TCR^+^ and γδ TCR^+^ subset definitions. **C**. Frequency of γδ TCR^+^ Vδ2^+^ T-cells among CD3^+^ T-cells following 12 days of stimulation with CBM588 (10 mg/mL) or HMBPP (10 ng/mL) in healthy donors (n = 28). Squares indicate donors in Japan, and circles indicate donors in the United Kingdom. Two-tailed Wilcoxon matched-pairs signed-rank test (**** p < 0.0001) and (* p = 0.0294). Box plot showing the median, and whiskers at minimum and maximum values.

Encouraged by these findings, we tested PBMCs from four additional healthy donors to confirm that the observed expansion of Vδ2⁺ T-cells was attributable to *Clostridium butyricum* spores rather than tablet excipients. As a positive control, we included (*E*)-4-Hydroxy-3-methyl-but-2-enyl pyrophosphate (HMBPP), a microbial metabolite from the non-mevalonate isoprenoid biosynthesis pathway (13). HMBPP activates Vγ9Vδ2 T-cells by acting as a molecular ’glue’ that promotes the heteromeric association of butyrophilins BTN3A1 and BTN2A1, forming a TCR ligand (14, 15). Robust induction of Vδ2⁺ T-cells was observed following stimulation with either the CBM588 tablet suspension, the purified spore preparation, or HMBPP (**Fig. 1B; Supplementary Fig. 2A**), with no expansion of other T-cell subsets (**Supplementary Fig. 2B**).

Combined priming experiments using PBMCs from 28 donors in the UK and Japan confirmed that CBM588 tablet suspension significantly expanded Vδ2⁺ T-cells after 12 days, although the magnitude of expansion varied widely (**Fig. 1C**). Notably, Vδ2⁺ T-cell frequencies were generally higher in the donors from Japan, with ∼30% of UK donors showing no expansion. This difference may reflect methodological or population-specific factors, or the fact that CBM588 is widely used in Japan to treat diarrhoea (16), potentially stimulating T-cell memory responses. Further investigation will be required to clarify this observation. Together, these data indicate that CBM588-derived bacterial components can expand Vγ9Vδ2⁺ T-cells from healthy donor PBMCs. As some Vγ9Vδ2⁺ T-cells are known to kill cancer cells in response to tumour-derived pyrophosphates (17–20), we next investigated whether CBM588-induced T-cells could respond to cancer cells.

### CBM588-primed γδ T-cells recognize cancer cells via BTN3A

To assess whether CBM588-primed γδ T-cells could recognize cancer cells, we used the lung cancer cell line A549 and generated a BTN3A knockout variant (A549-BTN3A KO) by CRISPR/Cas9 disruption of all three BTN3A isoforms, as described in the Materials and Methods. BTN3A1 is required for recognition of phosphoantigens such as HMBPP and isopentenyl pyrophosphate (IPP) by Vγ9Vδ2 T-cells (14), enabling us to test whether recognition was mediated via tumor-derived mevalonate metabolites (18). We assessed T-cell responses by flow cytometry at the single-cell level, measuring CD107a (a surrogate for degranulation and cytotoxic activity (21)) and surface TNF (**Supplemental Fig. 3**). In the UK-based cohort, 2 of 11 donors showed baseline reactivity to CBM588, and only one donor exhibited recognition of A549 lung cancer cells without priming. Following CBM588 priming, all 11 donors responded to CBM588, and 7 of 11 recognized A549 lung cancer cells (**Fig. 2A**). This recognition was abrogated in A549-BTN3A KO cells, consistent with a BTN3A-dependent mechanism involving IPP accumulation (18). While priming with HMBPP successfully induced CBM588-reactive T-cells, it did not confer cancer cell recognition (except in one donor with pre-existing reactivity) highlighting a key distinction between HMBPP and CBM588 (**Fig. 2A**). Both CBM588 and HMBPP-primed T-cells responded robustly to HMBPP, but only CBM588 priming induced cancer-reactive Vγ9Vδ2 T-cells, suggesting differential induction of TCR clonotypes or sensitivity. We then extended the analysis to include donors in Japan, combining data across 24 healthy individuals. CBM588-primed T-cells consistently recognised A549 cells via a BTN3A-dependent mechanism (**Fig. 2B**; **** p < 0.0001). In contrast, HMBPP priming did not increase A549 reactivity above baseline (**Fig. 2B**). Importantly, γδ TCR negative T-cells showed no response to A549 lung cancer cells under any priming condition (**Supplemental Fig. 3C**). Together, these findings indicate that CBM588 can prime Vγ9Vδ2 T-cells with the capacity to recognise tumor cells through a BTN3A-dependent mechanism. We next asked whether the CBM588-primed T-cells could kill lung cancer cells.

**Figure 2:**
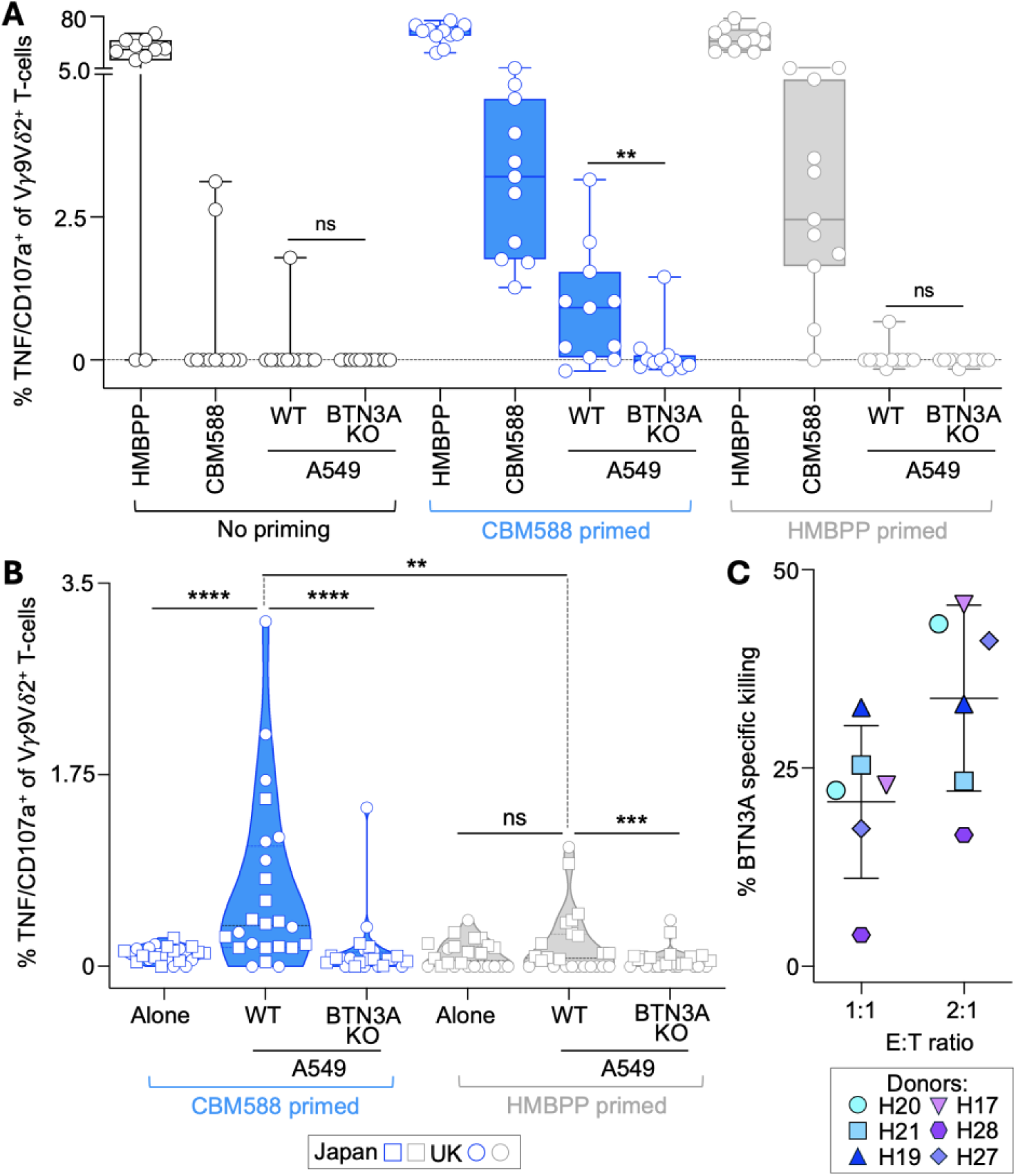
Cancer reactivity of CBM588 tablet-primed Vγ9Vδ2 T-cells from healthy donors. **A**. PBMCs from 11 healthy donors in the United Kingdom were left unprimed or primed with CBM588 tablet suspension (10 mg/mL) or HMBPP (10 ng/mL). After 12 days in culture, Vγ9Vδ2⁺ T-cell reactivity was assessed by T107 assay (CD107a/TNF) in response to CBM588 tablet suspension, HMBPP, lung cancer A549 wild-type or BTN3A knockout (KO) cells. Background levels of CD107a/TNF from unstimulated T-cells were subtracted. Data is presented as a box plot with median values and whiskers indicating minimum and maximum values. Two-tailed Wilcoxon matched-pairs signed rank test (** p = 0.0039). See **Supplementary Fig. 3A** for flow cytometry gating strategy. **B**. Vγ9Vδ2⁺ T-cell reactivity (T107 assay) from CBM588- or HMBPP-primed PBMCs from healthy donors in the United Kingdom (same as in a, without background subtraction) and Japan (as indicated in the key) in response to A549 and A459-BTN3A KO cells. Two-tailed Wilcoxon matched-pairs signed rank test (** p = 0.0051; *** p = 0.0002; **** p < 0.0001)). See **Supplementary Fig. 3** for αβ TCR⁺ subset reactivity. **C**. Cytotoxicity assay of CBM588-primed T-cells from six healthy donors against A549 or A549-BTN3A KO cells over 72 hours. Data shows BTN3A-dependent killing, calculated by subtracting the killing of A549-BTN3A KO cells from that of wildtype A549 cells. See **Supplementary Fig. 4** for the gating strategy and recognition of additional cancer cell types (leukemia and pancreatic).

### CBM588-primed T-cells can kill cancer cell lines in a BTN3A-dependent manner and recognize different cancer types

To confirm the cytotoxic potential of CBM588-primed T-cells, we performed a flow cytometry-based killing assay as previously described (22) (gating strategy: **Supplemental Fig. 4A**). In six healthy donors tested, CBM588-primed T-cells demonstrated BTN3A-dependent cytotoxicity against A549 lung cancer cells (**Fig. 2C**), reinforcing that CBM588 priming can elicit functionally potent Vγ9Vδ2 T-cells capable of tumor cell lysis via a BTN3A-dependent mechanism. We explored whether this effect extended beyond lung cancer. CBM588 exposure enhanced the responsiveness of donor γδ T-cells against additional cancer cell types, including pancreatic adenocarcinoma and leukaemia (**Supplementary Fig. 4B**), suggesting a broader cancer-recognition potential. We next investigated the impact of CBM588 on γδ T-cell responses in lung cancer patients receiving immune checkpoint inhibitors (ICIs), with or without concurrent administration of CBM588.

### Vγ9Vδ2 T-cells in lung cancer patients treated with ICIs and CBM588 exhibit increased activation and infiltration into cancerous site

To investigate the frequency and activation status of Vγ9Vδ2^+^ T-cells in lung cancer patients undergoing treatment with ICIs, we analyzed PBMCs from NSCLC patients who had been enrolled in trials of CBM588 therapy (patient details in **Fig. 3A** and **Supplementary Fig. 5**). For 28 patients, paired PBMC samples pre- and post-treatment with ICI (n = 13) or ICI+CBM588 (n = 15) were available for comparison, which showed no increase in the proportion of various αβ TCR^+^ and γδ TCR^+^ T-cell subsets once therapy had commenced, including Vδ2^+^ T-cells (**Supplementary Figure 6**). Next, we examined various activation markers on peripheral blood T-cells subsets and compared their pre- and post-therapy levels between the ICI (n = 13) and ICI+CBM588 (n = 22) patient cohorts. There was significant increase (p = 0.043) in the proportion of post-therapy CD69^+^ Vδ2^+^ T-cells in patients using ICI+CBM588 compared to ICI alone, suggesting *in vivo* activation (**Fig. 3B** and **Supplementary Fig. 7**). In contrast, Vδ1^+^ T-cells did not show any differences in CD69 expression between the ICI and ICI+CBM588 cohorts (**Supplementary Figs. 7 and 8A**). We also observed a significant increase (p= 0.0041) in the percentage of CD69^+^Vδ2^+^ T-cells between pre- and post-therapy samples for patients taking CBM588, but not for the ICI only cohort (**Fig. 3C** and **Supplementary Figs. 7 and 8B**). PBMCs from these patients, when primed with CBM588 for 12 days, were capable of recognizing (**Fig. 3D**) and killing (**Fig. 3E**) A549 lung cancer cells in a BTN3A-dependent manner. We also examined malignant pleural effusions from lung cancer patients treated with ICI alone or in combination with CBM588. The proportion of Vγ9Vδ2 T-cells was significantly higher in patients receiving ICI+CBM588 compared to those on ICI alone (**Fig. 3F**). This increase in Vγ9Vδ2 cells was accompanied by a relative decrease in Vδ1^+^ T-cells (**Fig. 3F**), and no αβ TCR^+^ subsets were altered (**Supplementary Fig. 9A**), thereby indicating that CBM588 alters the γδ T-cell niche at tumor sites. Overall, these data demonstrate increased activation and intratumoral infiltration of Vγ9Vδ2 T-cells in lung cancer patients treated with CBM588, providing a plausible mechanistic basis for the improved patient outcomes observed across multiple clinical studies (9–12).

**Figure 3:**
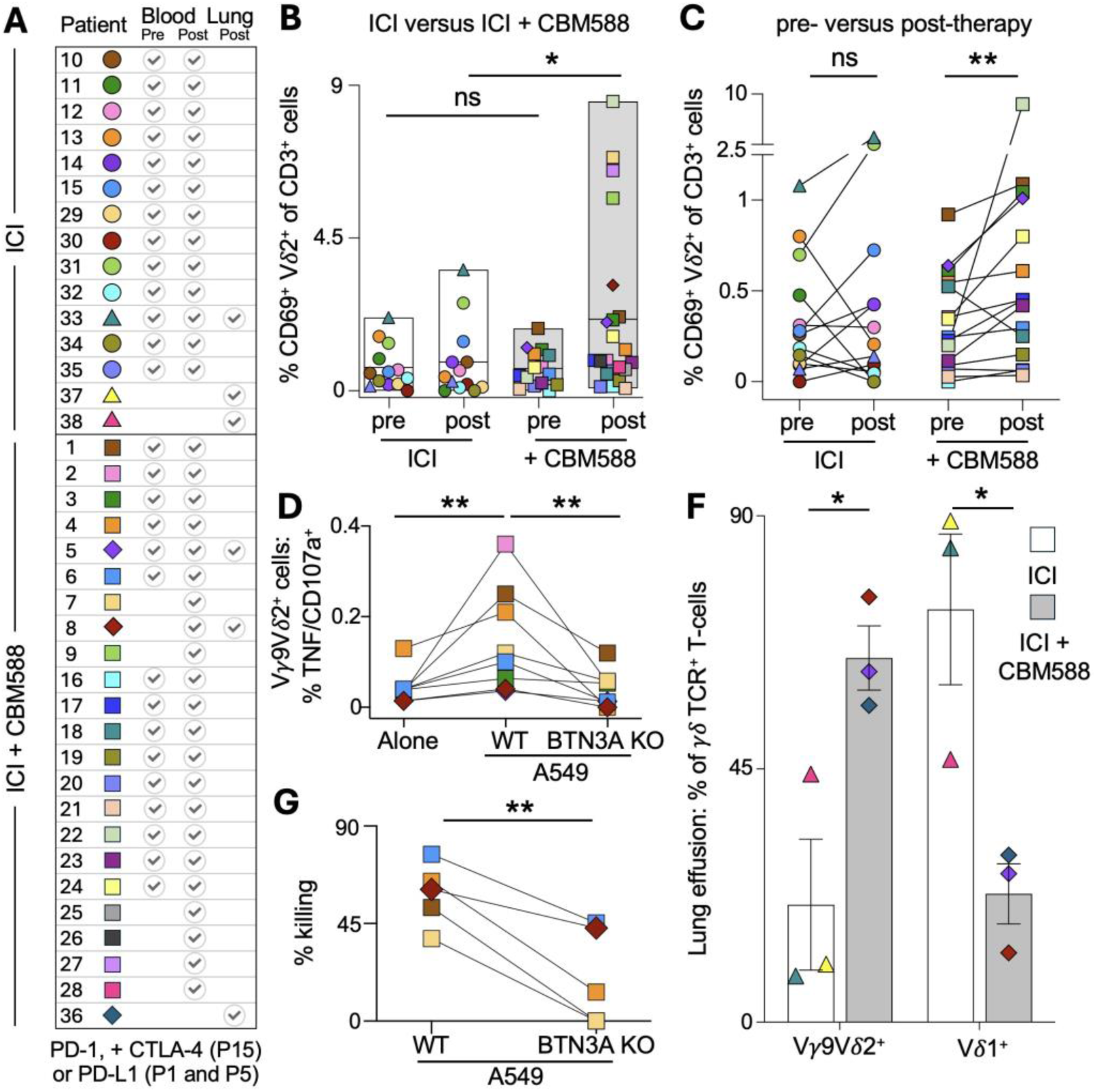
Frequency, activation status and cancer reactivity of Vδ2^+^ T-cells in lung cancer patients treated with ICI and CBM588. **A**. Lung cancer patients based in Japan received immune checkpoint inhibitor (ICI) therapy with or without CBM588. Each patient is represented by a unique symbol and color combination. PBMCs were collected pre- and/or post-treatment; pleural effusion samples were collected post-treatment from six patients. See **Supplementary Fig. 5** for patient demographics and sampling details. **B**. Comparison of CD69^+^ Vδ2^+^ T-cell frequency between ICI and ICI+CBM588 treatment groups using all patients with PBMC samples (n = 35). Two-tailed Mann– Whitney test (* p = 0.043). **C**. Frequency of CD69^+^ Vδ2^+^ T-cells within CD3^+^ T-cells from patients with available pre- and post-treatment PBMC samples (n = 28). Two-tailed Wilcoxon matched-pairs signed rank test (** p = 0.0041). See **Supplementary Figures 7&8** for expression of other activation markers and T-cell subsets. **D**. PBMCs from patients treated with ICI+CBM588 were primed *in vitro* with CBM588 tablets and assessed for cancer reactivity using the T107 assay (CD107a/TNF) against A549 and A549-BTN3A KO lung cancer cells. Two-tailed Wilcoxon matched-pairs signed rank test (** p = 0.0078). **E**. Killing assay using CBM588-primed PBMCs from patients treated with ICI+CBM588, challenged with A549 or A549-BTN3A KO cells (1:1 ratio, 72 h). Paired two-tailed Student’s t-test (** p = 0.0040). **F**. Proportion of γδ TCR^+^ T-cell subsets in pleural effusions from lung cancer patients treated with ICI with or without CBM588. Error bars represent standard error of the mean. See **Supplementary Fig 9** for additional T-cell subset data from pleural effusions. Paired two-tailed Student’s t-test (* p = 0.0276) for Vγ9Vδ2^+^ and (* p = 0.0248) for Vδ1^+^ T cells.

### Activation of Vδ2^+^ T-cells in lung cancer patients correlates with improved outcomes

We next assessed the relationship between Vδ2^+^ T-cell activation and clinical outcomes. To avoid bias, we analyzed the ICI and ICI+CBM588 arms as a single cohort, given that CBM588 is already known to significantly improve outcomes (9, 10). The median frequency of CD69 expression on Vδ2^+^ T-cells was calculated, and patients were stratified into two groups: those with CD69 expression above the median and those with expression equal to or below it, both pre- and post-treatment (**Supplementary Fig. 9B**). CD69 expression before treatment did not correlate with overall survival, but post-treatment expression was strongly predictive (p < 0.03) (**Figs. 4A&B**). All patients in the “CD69 low” group succumbed to disease, whereas over half of the “CD69 high” group were still alive at the time of being monitored post treatment (0.81 to 7.12 years, with a mean of 3.04 years; **Figure 4B**). Most of the long-term survivors had stable disease, and all three patients who achieved complete remission were in the CD69 high group (**Figure 4C**) and had received CBM588. Fewer than half of patients were CD69 high before treatment with ICI+CBM588; this proportion rose to nearly three-quarters post-treatment (**Figure 4D**). Significantly improved survival and clinical outcomes were also observed when Vδ2^+^ T-cell analysis was based on an increase in CD69 expression following the initiation of therapy (**Supplementary Fig. 10**).

**Figure 4.**
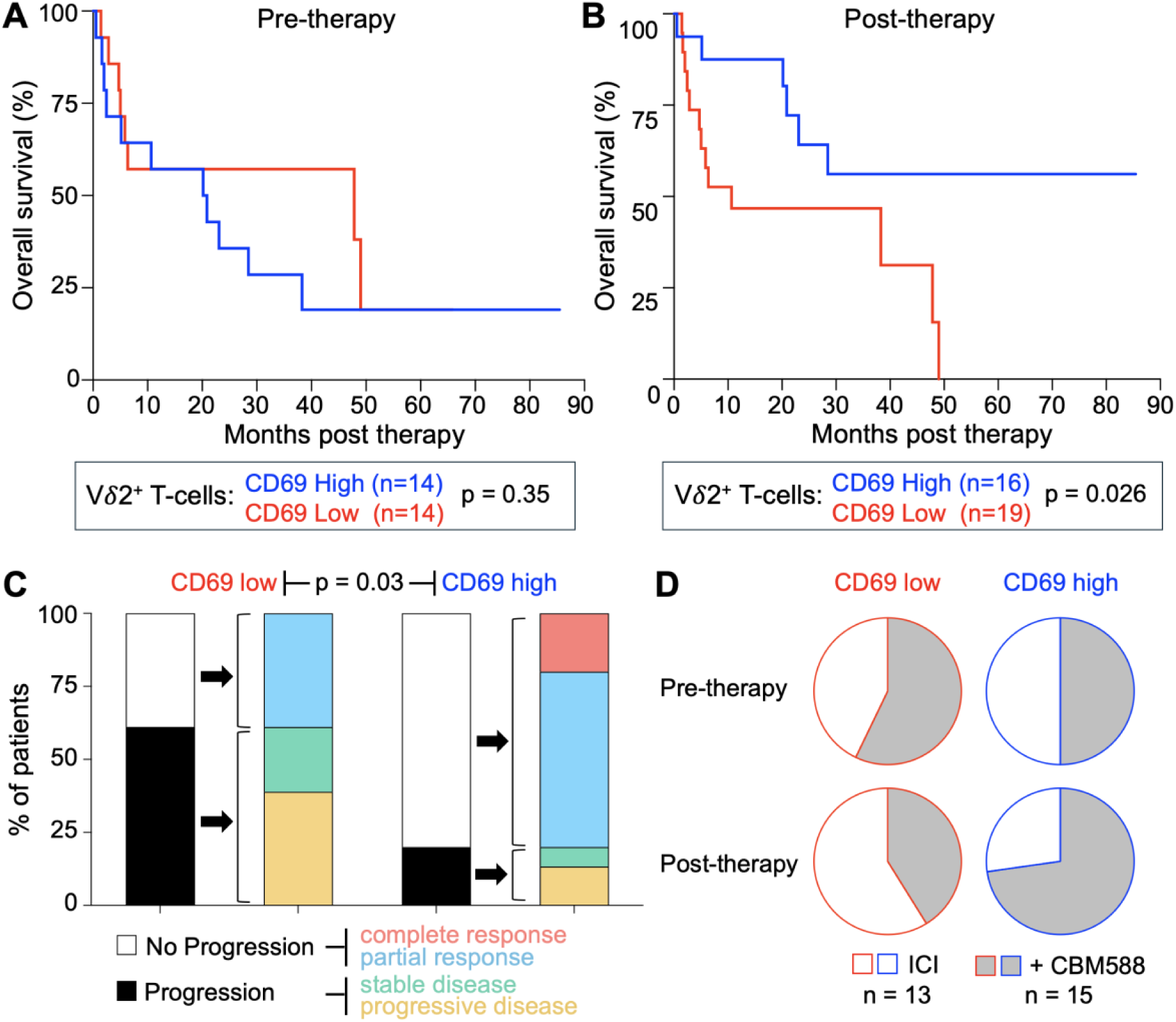
High CD69 expression on Vδ2⁺ T-cells correlates with improved outcomes in lung cancer patients receiving immunotherapy. **A**. Kaplan–Meier analysis of overall survival in lung cancer patients (n = 28, with pre- and post-therapy samples) treated with ICI or ICI+CBM588 (analysed as a combined cohort). Patients were stratified into high or low CD69 expression groups based on CD69^+^Vδ2^+^ T-cell frequency in pre-treatment blood samples, relative to the cohort median. *p* value from a log-rank test. Median values of CD69 are shown in **Supplementary Fig 9**. **B**. Kaplan– Meier analysis as in (**a**), based on CD69⁺Vδ2⁺ T-cell frequency in post-treatment blood samples from patients (n = 35, with post-therapy samples). *p* value from a log-rank test. **C**. Clinical outcomes of patients (n = 33, clinical response category nonevaluable for two patients) stratified by high or low CD69 expression on Vδ2^+^ T-cells post-therapy. Outcomes categorized according to the key. Statistical analysis by a Fisher’s exact test. **D**. Distribution of patients (n = 28, with pre- and post-therapy samples) treated with ICI or ICI+CBM588 across the pre- and post-treatment CD69 high/low Vδ2⁺ T-cell categories. See **Supplementary Fig 10** for related analyses based on post-therapy increases or decreases in CD69 expression.

## DISCUSSION

Recent findings suggest that the gut microbiome plays a key role in cancer prognosis and response to treatment. Gajewski and colleagues showed that melanoma growth varied markedly in mice populated with distinct commensal microbiota, with *Bifidobacterium* associated with antitumour effects (7). Oral administration of *Bifidobacterium* was as effective as checkpoint blockade in these animals, with combination treatment almost eliminating tumour outgrowth (7). In humans, the composition of the gut microbiome differs significantly between responders and non-responders to anti-PD1 immunotherapy in metastatic melanoma (5, 6). Clinical trials have also shown that faecal microbiota transplantation can influence the response in patients with anti-PD1-refractory melanoma (23–25). A further study linked specific microbial species in the baseline stool microbiome to improved response rates and prolonged progression-free survival in patients with epithelial tumors (4). Several recent studies have shown that gut microbiome composition serves as a predictive biomarker for response to CD19-targeted CAR-T cell therapy (26, 27), with one particularly striking report demonstrating that supplementation with the newly classified bacterial species *Akkermansia massiliensis* enhances the therapeutic potency of CAR-T cells against B-cell malignancies (28). These findings across multiple malignancies have generated considerable interest in whether probiotics could be harnessed to improve treatment outcomes and cancer survival.

Optimal exploitation of the intestinal microflora in cancer therapy will require a mechanistic understanding of how gut microorganisms exert effects at distant tumor sites. Several possibilities have been proposed, including direct or indirect modulation of the tumor-resident microbiome, alteration of the immune microenvironment, and the systemic effects of microbial metabolites on immune and/or tumor cells. Notably, human unconventional T-cells, including γδ T-cells, can respond to intermediates in microbial metabolic pathways such as those for vitamin B and isoprenoid biosynthesis (18, 29), providing a direct route by which the microbiota can influence systemic T-cell activity. We were interested in whether a probiotic species associated with improved cancer outcomes might be capable of directly priming T-cells that recognize and respond to cancer cells.

Several recent studies have shown that daily administration of CBM588 significantly improves response rates and overall survival in patients with NSCLC and mRCC when used alongside ICIs (9–12). The mechanisms underlying these clinical benefits remain unclear, and only a subset of patients appear to derive full advantage from treatment. In the study by Dizman and colleagues (11), wide differences in metabolic pathways emerged between CBM588-treated and untreated patients over a 12-week period. Here, we show that *in vitro* stimulation of PBMC with CBM588 can stimulate the expansion of a population of cancer-reactive Vγ9Vδ2 T-cells with the capacity to kill cancer cell lines via a BTN3A-dependent mechanism. However, there was substantial variability between donors. Nearly a third of healthy donors showed little or no expansion of Vγ9Vδ2 T-cells in response to CBM588, and in some cases, expanded cells lacked reactivity against cancer cell lines. This variability suggests important differences in TCR specificity or sensitivity to antigen, and further work will be needed to understand how this might relate to differential patient responses to probiotic therapy. Importantly, using PBMCs from lung cancer patients in recent studies conducted in Kumamoto (9, 10), we found that the frequency of activated (CD69⁺) Vδ2⁺ T-cells in peripheral blood increased following CBM588 treatment. In patients receiving ICI+CBM588, the proportion of CD69⁺Vδ2⁺ T-cells was significantly higher than in those receiving ICI alone, suggesting *in vivo* activation of these cells. Notably, elevated levels of CD69 expression on in peripheral blood Vδ2⁺ T-cells post-therapy correlated with improved clinical outcomes and overall survival in lung cancer patients. CBM588 also influenced the γδ T-cell landscape within diseased tissue; lung effusions from patients receiving ICI+CBM588 contained a significantly greater proportion of Vγ9Vδ2 T-cells compared to patients on ICI alone. These findings suggest that CBM588 acts to promote the expansion and activation of cancer-reactive Vγ9Vδ2 T-cells at the cancer site in humans.

Overall, our results support a model in which certain gut-resident bacteria, such as *Clostridium butyricum*, can influence systemic antitumour immunity through selective engagement of the γδ T-cell compartment. These findings open new avenues for understanding and harnessing microbial control of cancer immunosurveillance. In particular, microbiota-derived isoprenoids warrant further investigation. Isoprenoids and their derivatives constitute the largest family of natural organic compounds, with over 65,000 members found across all domains of life (30, 31). Two evolutionarily distinct biosynthetic pathways exist for their production (32), and organisms may employ either or both. The diversity of enzymes and intermediates, especially in the early steps, continues to expand (33). Mammalian cells utilize the mevalonate (MVA) pathway, which produces isopentenyl pyrophosphate (IPP), a known ligand for Vγ9Vδ2 T-cells (18). Most bacteria, by contrast, use the non-mevalonate (MEP) pathway, which generates ligands such as HMBPP that are vastly more potent than IPP in stimulating these cells. Remarkably, IPP and HMBPP differ by only a single hydroxyl group, yet HMBPP is ∼10,000-fold more effective at activating cognate T-cells (34).

Our findings demonstrate that CBM588 can directly induce cancer-reactive Vγ9Vδ2 T-cells and remodel the γδ T-cell landscape in both peripheral blood and the tumor site, providing a plausible immunological mechanism for its clinical benefit. We also identify Vδ2⁺CD69⁺ T-cells in patient blood as a robust prognostic marker in NSCLC. These observations are consistent with a comprehensive pan-cancer analysis of over 18,000 human tumors, which identified γδ T-cell infiltration as the strongest predictor of favorable prognosis across 25 malignancies, exceeding the prognostic value of αβ T-cell subsets (35). Taken together, these results reveal the gut microbiome as a key modulator of systemic antitumour immunity and suggest that its rational manipulation could reshape the immunological landscape of cancer. Probiotic interventions, such as CBM588, may ultimately offer a tractable route to enhance therapeutic efficacy and improve clinical outcomes across diverse malignancies.

### Study limitations

CBM588 has been shown to improve clinical outcomes in NSCLC and RCC, but the mechanisms underlying these effects have remained unclear. In this study, we provide evidence that CBM588 can expand cancer-reactive Vδ2⁺ T-cells from the PBMC of both healthy donors and patients with lung cancer. In patients initiating immune checkpoint blockade alongside CBM588, we also observe marked shifts in the peripheral and tumor-associated γδ T-cell compartment, as well as a correlation between Vδ2⁺ T-cell activation and improved survival. However, several limitations should be acknowledged. First, our study is observational and relies on samples obtained from retrospective clinical studies, which limits the ability to draw definitive causal inferences. While our data suggest that CBM588 supplementation modulates the Vδ2⁺ T-cell compartment *in vivo*, we cannot rule out the contribution of other factors associated with treatment or patient heterogeneity. Second, CBM588 is known to induce broad changes in the gut microbiome. As such, the immunological effects observed in patients could result not only from direct action of *C. butyricum*, but also from secondary shifts in microbiota composition, either through expansion of other beneficial species or suppression of inhibitory taxa. Finally, although Vδ2⁺ T-cell activation correlated with improved outcomes in our patient cohort, this association does not prove mechanistic linkage. It remains possible that Vδ2⁺ activation reflects an indirect consequence of improved clinical outcome rather than a driver of therapeutic benefit. Notably, a recent study has shown that certain commensal bacteria can augment checkpoint blockade by driving the migration of activated dendritic cells from the gut to the tumor site, thereby enhancing CD8⁺ T-cell priming and infiltration (36). Such findings highlight the likelihood that multiple immunological mechanisms may operate in parallel to mediate microbiome-driven cancer control. Our results provide a plausible immunological mechanism for the clinical effects of CBM588 and offer a new framework for exploring microbiome–immune system interactions in cancer. Prospective studies will be essential to validate these associations, clarify causality, and establish whether Vδ2⁺ T-cells contribute directly to improved tumor control in patients receiving CBM588.

## METHODS

### Sex as a biological variable

Sex was not considered as a biological variable in this study.

### Patients

38 patients with NSCLC who received ICIs alone or ICIs + CMB588 (a dose of 120 mg orally three times daily) at Kumamoto University Hospital between 2017 and 2023 were enrolled in this study for flow cytometric analyses *ex vivo* and *in vitro* (approval number 1825). All human subjects provided written informed consent.

### Isolation of PBMCs from whole blood

Blood samples from healthy donors were sourced from the Welsh Blood Service (Velindre NHS Trust, Wales, UK) as EDTA treated ‘buffy coats’ and ethical approval granted by the School of Medicine Research Ethics Committee (reference 18/56). Each buffy coat was seronegative for HIV-1, HBV and HCV. Blood and cells derived thereof were handled in accordance with Cardiff University guidelines in alignment with the United Kingdom Human Culture Act 2004. Blood from the WBS was diluted 1:1 with R10 (RPMI-1640 supplemented with 10% heat-inactivated FBS, 100 U/mL penicillin, 100 µg/mL streptomycin and 2 mM L-glutamine (all Merck, New Jersey, US)) and placed on a Cole-Parmer™ Stuart™ roller-mixer overnight at 9-11 rpm and room temperature. The following morning the blood was further diluted 2:1 (blood:RPMI-1640) then PBMCs separated using conventional density gradient centrifugation with Sigma-Aldrich Histopaque 1077 (Merck). Red blood cells were removed using lysis buffer (155 mM NH4Cl, 10 mM KHCO3 and 0.1 mM EDTA, pH7.2-7.4) for 10 min at 37°C. PBMCs were used fresh for culture, without cryopreservation. Blood samples were also obtained from Japanese healthy volunteers. The blood diluted 2:1 (blood:RPMI-1640) and then PBMCs were purified by a density gradient centrifugation using Ficoll-Paque Plus (GE Healthcare Life Sciences, Cat# 17-1440-03) and stored in liquid nitrogen until further use.

### Cell Culture

Cells were cultured at 37°C and 5% CO_2_ and were tested once a month for mycoplasma using a MycoAlert^®^ mycoplasma detection kit according to the manufacturer’s instructions (Lonza, Basel, Switzerland). Cell lines were validated based on morphology, characteristics, and phenotypic markers according to the American Type Culture Collection (ATCC) or the German Collection of Microorganisms and Cell Cultures (Leibniz Institute DSMZ).

### Cancer Cell Lines

The following cell lines were used (reference for cell line information included in brackets): lung carcinoma A549 (ATCC® CCL-185™), chronic Myelogenous Leukemia K562 (ATCC® CRL-3344™) and pancreatic adenocarcinoma DAN-G (DSMZ ACC249). K562 cells were grown in R10 as suspension cultures and split 1:10 once or twice a week. A549 (R10 media) and DAN-G (R10 media) cells lines are adherent, and were passaged by detachment with D-PBS + 2 mM EDTA and split 1:10-1:20 once a week. A549 were engineered for the study as described below, to remove expression of butyrophilin 3A family members.

### T-cell lines

PBMCs were primed with CBM588 tablet or spores (Miyarisan Pharmaceutical Co. Ltd. Tokyo, Japan), or (*E*)-1-Hydroxy-2-methyl-2-butenyl 4-pyrophosphate lithium salt (HMBPP) (Merck) at concentrations indicated and cultured in priming media (RPMI-1640, 100 U/mL Penicillin, 100 µg/mL Streptomycin, 2 mM L-Glutamine, 1X non-essential amino acids, 1 mM sodium pyruvate, 10 mM HEPES buffer, 10% FBS (Gibco, MA, USA) and 20 IU/mL IL-2 (Aldeslukin, Proleukin; Prometheus, San Diego, CA, USA) for two weeks.

### Generating A549 butyrophilin 3A knockout cells

A549 BTN3A knockout cells were generated using a CRISPR/Cas9 lentiviral system based on the on the lentiCRISPR v2 plasmid (Addgene plasmid #52961, a kind gift from Feng Zhang). The puromycin selection gene was replaced with truncated (t) NGFR (CD271) by the Gibson Assembly technique to aid selection of knockout cells(37). The gRNA (5’-ATCAGCGTCTTCACCCACCA TGG-3’) targeting all three BTN3A isoforms (BTN3A1, A2 and A3) was cloned into the lentiCRISPR v2-tNGFR plasmid as described by Addgene. The knockout plasmid was transformed into XL10 gold competent cells (Agilent, CA, USA) for amplification and plasmid Minipreps performed using PureLink™ Quick Plasmid Kit (Life Technologies, CA, USA). For transfection, the lentiCRISPR v2-BTN3A KO gRNA-tNGFR plasmid, the pMD2.G envelope plasmid (Addgene plasmid #12259) and the psPAX2 packaging plasmid (Addgene plasmid #12260) in Opti-MEM (Gibco) media with the addition of 1µg/µL polyethylenimine (Merck) were incubated at room temperature for 15 min then added to HEK293T cells that were cultured in D10 media (as for R10, but with DMEM (Merck)) and incubated at 37°C with 5% CO_2_ for three days, with media being removed and replaced with fresh D10 each day. The superna:t media from 48 and 72 hours post-transfection were kept, combined, centrifuged at 800 x g for 5 min and filtered using a 0.45 µM filter (Fisher Scientific, MA, USA). A549 cells were plated at 100,000 cells per 24-well in 1 mL of R10 media and cultured overnight. The next day media was replaced with 500 µL fresh R10 and 500 µL lentivirus supernatant with 5 µg/mL polybrene (Sigma Aldrich, MO, USA) and cells were centrifuged at 400 x g for 2 h for spinfection. Cells were cultured for 7 days then tested for BTN3A knockout compared to the A549 wild-type cell line by antibody staining towards co-marker CD271/tNGFR (clone ME20.4-1.H 4, Miltenyi Biotec) and CD277/BTN3A (clone BT3.1, also known as clone 20.1, Miltenyi Biotec) surface antibodies (CD277 antibody required a 45 min at 4°C; standard 20 min incubation on ice used for all other surface antibodies). The A549 BTN3A knockout cell line was then cloned by limiting dilution and clones were confirmed for knockout by surface antibody staining, genomic sequencing to reveal insertion or deletions, and functional testing with γ9δ2 T-cell lines following pretreatment with 50 µM Zoledronate (Sigma Aldrich).

### In vitro induction of CBM588 or HMBPP-reactive T cells from PBMCs

A suspension of CBM588 tablets was made by dissolving the tablets in PBS (100 mg/mL) and vortexing it for 1 min. Human PBMCs from healthy donors were incubated with 2.5, 5 or 10 mg/mL of CBM588 suspension or 1, 10 or 100 ng/mL of HMBPP in 96U wells (1 x10^6^ PBMCs per well) with 200 µL per well of priming media for 12 days, with the 50% of the media being changed two times per week. Post optimisation, the standard concentrations of CBM588 tablet or HMBPP used for priming T-cells was 10 mg/mL and 10 ng/mL respectively.

### Phenotyping of PBMCs primed with CBM588 or HMBPP

*In vitro* primed T cells were washed and stained with Fixable Live/Dead Violet Dye (1/40 pre-dilution then 1/50 for staining; , and surface stained with following antibodies: For the antibody panel used in the UK, CD4 BV510 (SK3, 1/100 dilution: BioLegend, San Diego, CA, USA), Vδ2 PE (REA771, 1/100 dilution; Miltenyi Biotec), CD3 PerCP (UCHT1, 1/100 dilution; BioLegend), γδ TCR APC (REA591, 1/100 dilution; Miltenyi Biotec), Vγ9 APC Vio770 (REA470, 1/100 dilution; Miltenyi Biotec), CD8 PEvio770 (REA734, 1/200 dilution; Miltenyi Biotec), TCRαβ FITC (IP26, 1/100 dilution; BioLegend). For the panel used in Japan, Vδ2 BioBlue (REA771, 1/100 dilution; Miltenyi Biotec), Vδ1 PE-eFluor610 (TS8.2, 1/25 dilution; eBioscience), Vγ9 TCR APCvio770 (REA771, 1/100 dilution; Miltenyi Biotec), αβ TCR PerCP/Cy5.5 (IP26, 1/100 dilution; BioLegend), γδ TCR FITC (REA591, 1/100 dilution; Miltenyi Biotec), CD3 BV785 (UCHT1, 1/50 dilution; BioLegend), CD8 BV570 (RPA-T8, 1/100 dilution; BioLegend), CD4 BV750 (SK3, 1/100 dilution; BioLegend), CD69 BV421 (FN50, 1/100 dilution; BioLegend), PD-1 APC (MIH4, 1/50 dilution; BioLegend), CD137 BV711 (4B4-1, 1/50 dilution; BioLegend), CD103 PerCP-eFluor710 (Ber-ACT8, 1/25 dilution; eBioscience), CD38 AF647 (HIT2, 1/100 dilution; BioLegend), CD39 BV605 (A1, 1/50 dilution; BioLegend), CD56 BV510 (HCD56, 1/50 dilution; BioLegend), CD95 BV650 (DX2, 1/25 dilution; BioLegend), TIM3 PE (F38-2E2, 1/25 dilution; BioLegend), 7AAD (1/50 dilution; Biolegend), After incubation for 20 min on ice, the cells were fixed with 2% paraformaldehyde made in-house using formaldehyde or purchased commercially (Nacalai Tesque, Kyoto, Japan, Cat# 09154-85), and the levels of protein surface expression were analyzed by flow cytometry using a FACS Canto II (BD Biosciences, New Jersey, USA) or Cytek Northern Lights (Cytek Japan). The data obtained by flow cytometry were analyzed with FACS Diva v9.0 (BD) and FlowJo software v10 (Tree Star).

### T107 T-cell activation assay

Unprimed, CBM588- and HMBPP-primed T cells were rested in R5 (as for R10 with 5% FBS) for 24 h before assay. Typically, 1×10^5^ T cells and 2×10^5^ target cells 5 mg/mL CBM588 (per well), 1 μM HMBPP (262 ng/ml) or 2 μL CD3-CD28 Dynabeads™ (ThermoFisher) were co-incubated for 4 h with 30 μM TAPI-0 (cat# sc-203410, Santa Cruz Biotechnology, Texas, USA), and antibodies directed against TNF (cA2, PE Vio770, 1/100 dilution; Miltenyi Biotec) and CD107a (H4A3, FITC, 1/100 dilution; BD) or (H4A3, BV421, 1/100 dilution; BioLegend), were also included at the start of the assay (38). Following incubation, cells were washed and stained with Fixable Live/Dead Violet Dye, and the following conjugated antibodies: For antibody panel used in the UK, Vδ2 PE (REA771, 1/100 dilution; Miltenyi Biotec), CD3 PerCP (UCHT1, 1/100 dilution; BioLegend), αβ TCR BV510 (IP26, 1/100 dilution; BioLegend), γδ TCR APC (REA591, 1/100 dilution; Miltenyi Biotec), Vγ9 TCR APCvio770 (REA771, 1/100 dilution; Miltenyi Biotec). For the panel used in Japan, Vδ2 PE (REA771, 1/100 dilution; Miltenyi Biotec), CD3 BV750 (UCHT1, 1/50 dilution; BioLegend), αβ TCR PerCP/Cy5.5 (IP26, 1/100 dilution; BioLegend), γδ TCR FITC (REA591, 1/100 dilution; Miltenyi Biotec), CD8 BV570 (RPA-T8, 1/100 dilution; BioLegend), CD4 BV750 (SK3, 1/100 dilution; BioLegend) and levels of protein expression were analyzed by flow cytometry using a FACS Canto II (BD Biosciences) followed by analysis using FACS Diva v9.0 (BD) or Cytek Northern Lights (Cytek Japan) and FlowJo v10 software (Tree Star). Gating strategy in **Supplementary Fig. 3A and 3B**.

### Flow cytometry-based cancer cell killing assays

1×10^5^ target cell lines were plated in 96U-well plates, and CBM588-primed T cells added to give the desired T cell to target cell line ratio. Cells were co-cultured in 200 μL of target-cell media supplemented with 20 IU of IL-2 and 25 ng/ mL of IL-15 and incubated for 72 h. Before collection, 1×10^5^ A549-GFP cells were added to each well to allow the number of target cells that remained in experimental and control wells to quantified. Cells were washed three times with chilled D-PBS EDTA (2 mM) then stained in the assay plates with Fixable Live/Dead Violet Dye (VIVID, ThermoFisher), then CD3 APC (clone UCHT1, BioLegend) to allow dead cells and T-cells to be gated out, leaving viable target cells for analyses (**Supplementary Fig. 4A**). The data were analyzed by flow cytometry using a FACS Canto II (BD Biosciences) followed by analysis using FACS Diva v9.0 (BD) or Cytek Northern Lights (Cytek Japan) and FlowJo v10 software (Tree Star). Percentage killing was calculated as previously described (22).

#### Statistics and Reproducibility

Data and statistical analysis were performed using Prism 10 (GraphPad Software). For two-way comparison, the paired Wilcoxon signed-rank test (Fig. 1C, Fig. 2A, Fig. 2B, Fig. 3B, Fig. 3D, Fig. 3D, and Fig. 3E) and the paired T test (Fig. 3F) were used.

### Study Approval

For the use of human specimens, all protocols involving human subjects recruited at Cardiff University and Kumamoto University were reviewed and approved by the Institutional Review Boards. Donors recruited via the Welsh Blood Service gave informed consent as part of the donation procedure and samples used under local ethical approval granted by the School of Medicine Research Ethics Committee (reference 18/56). Donors recruited in Japan gave their consent and the study approved by Kumamoto University (approval number 1825).

### Data availability

The Supporting Data Values file contains all underlying values for data presented in this manuscript.

### Author Contributions

CM, YG and YT performed the experiments. TU, KI, TS and YT collected clinical samples. GD, HT, TM, AH, KO and MT prepared reagents. CM, YG, GD and AKS designed the experiments and interpreted the results. CM, GD and AKS wrote the original manuscript. All authors reviewed and proofread the manuscript. YG and GD share first authorship. The order of their names was agreed between them, with YG listed first in recognition of performing more of the experiments, and GD second in view of their primary role in data visualization and manuscript preparation.

## Acknowledgments

This work was supported in part by AMED Research Programs on HIV/AIDS (21fk0410046 to CM) and Interdisciplinary Cutting-edge Research (23wm0325064 to CM); JSPS KAKENHI Grants-in-Aid for Scientific Research B (22H02877 to CM), Scientific Research C (22K08256 to YT), and Fostering Joint International Research (A) (22KK0277 to CM); the Takeda Science Foundation (to CM); the Shinnihon Foundation of Advanced Medical Treatment Research (to YT); and the Kumamoto University Hospital Research Revitalization Project (to CM, TU, YT, and ST). AKS is a Wellcome Investigator (220295/Z/20/Z); GD and HT were funded through this award. This work was supported by Cancer Research Wales and TM was supported by the Cancer Research Wales Tom Walker Fund. We are deeply grateful to the Walker family for raising these funds in memory of Tom. We thank Philip Savage for critical reading of the manuscript.

