## Supplementart Figures for "A probiotic bacterium modulates antitumor γδ T-cell responses in lung cancer"

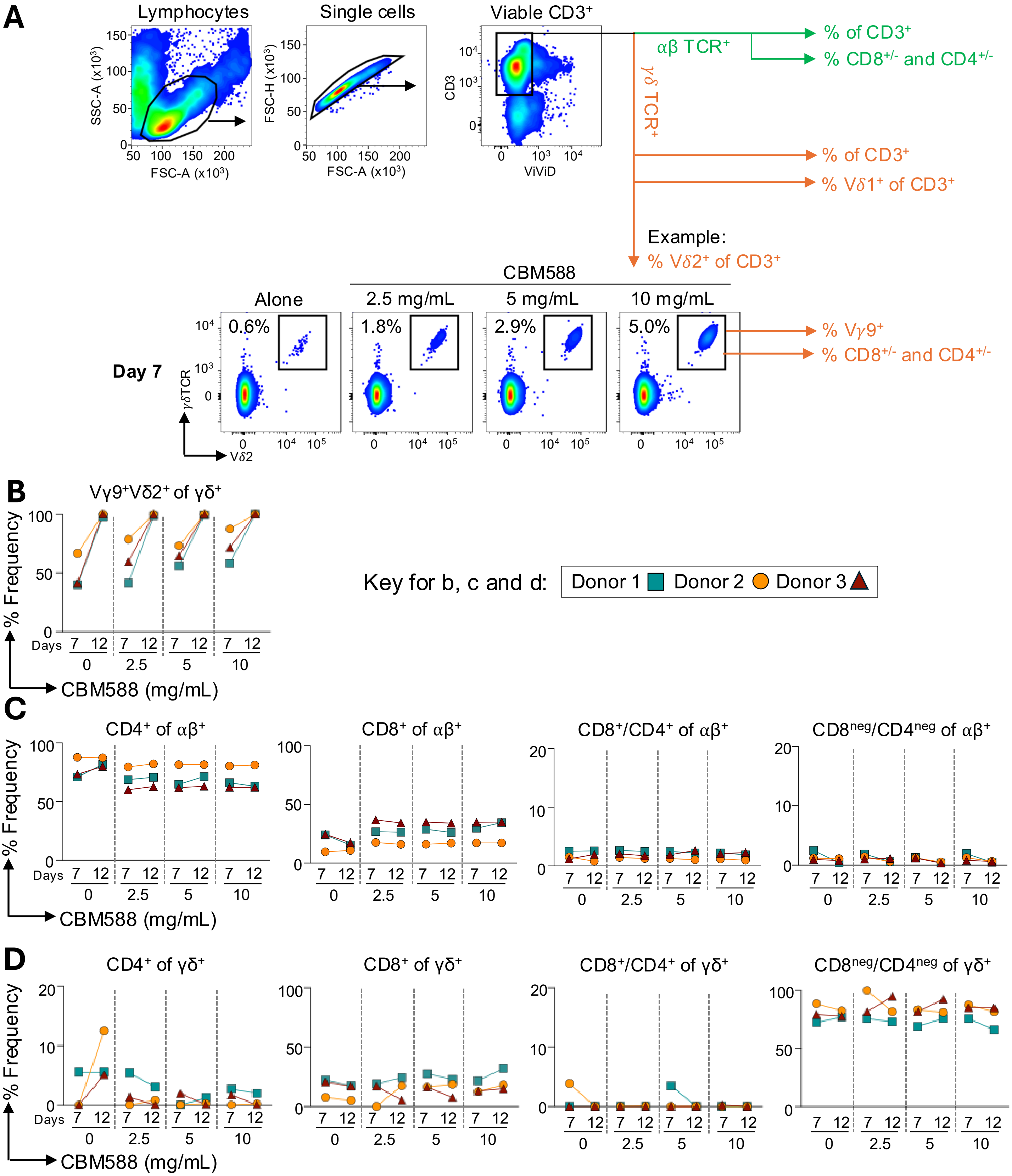


**Supplementary Figure 1. Flow cytometry gating strategies and T-cell dissection following priming with CBM588 tablet. A.** Flow cytometry gating strategies for different T-cell subsets following the priming of PBMCs with CBM588 tablet at different concentrations. Flow cytometry data for $\gamma\delta$ TCR⁺ V$\delta$2⁺ T-cells shown as an example for one of the donors. **B**. Frequency of $\gamma\delta$ TCR⁺ V$\gamma$9⁺ T-cells. **C**. Frequency of $\alpha\beta$ TCR⁺ T-cells, with analysis based on co-receptor expression. **D**. Co-receptor dissection of $\gamma\delta$ TCR⁺ T-cells.

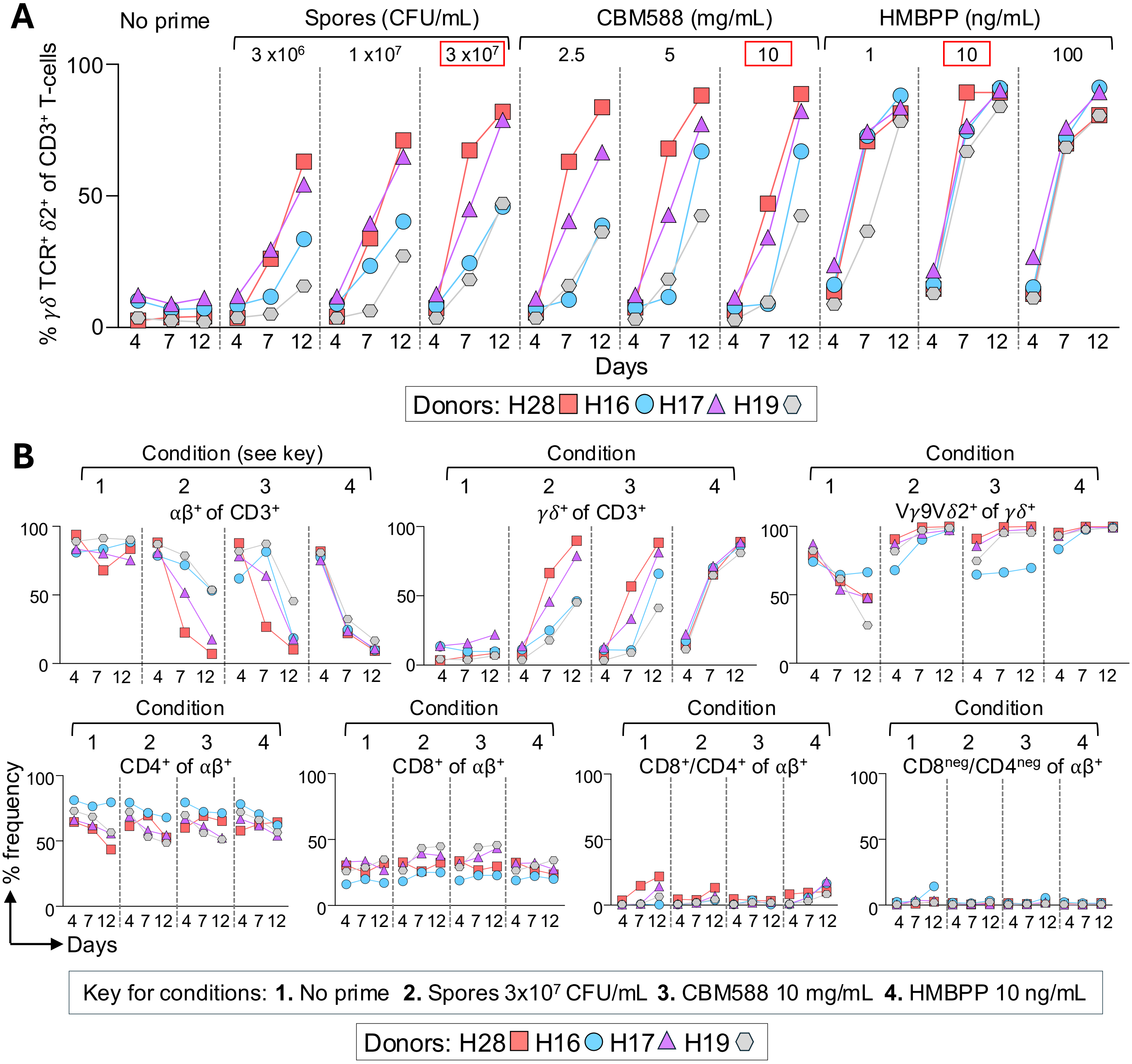


**Supplementary Figure 2. Frequency of $\gamma\delta$ TCR$^{+}$ V$\delta$2$^{+}$ T-cells following stimulation with CBM588 spore, CBM588 tablet and HMBPP. A.** Frequency of $\gamma\delta$ TCR$^{+}$ V$\delta$2$^{+}$ T-cells in CD3$^{+}$ T-cells after stimulation of PBMCs from four healthy donors with CBM588 spore, suspension of CBM588 tablet or HMBPP. T-cells sampled at days 4, 7 and 12 post stimulation. Data indicated by the red boxes, for colony forming units (CFU) of spore, and concentrations of CBM588 tablet and HMBPP, are also displayed in **Figure 1B**. **Bb** Frequency of T-cell subsets after stimulation of PBMCs for the four healthy donors in (a) with CBM588 spores, suspensions of CBM588 tablet, and HMBPP at the CFUs or concentrations indicated in the key.

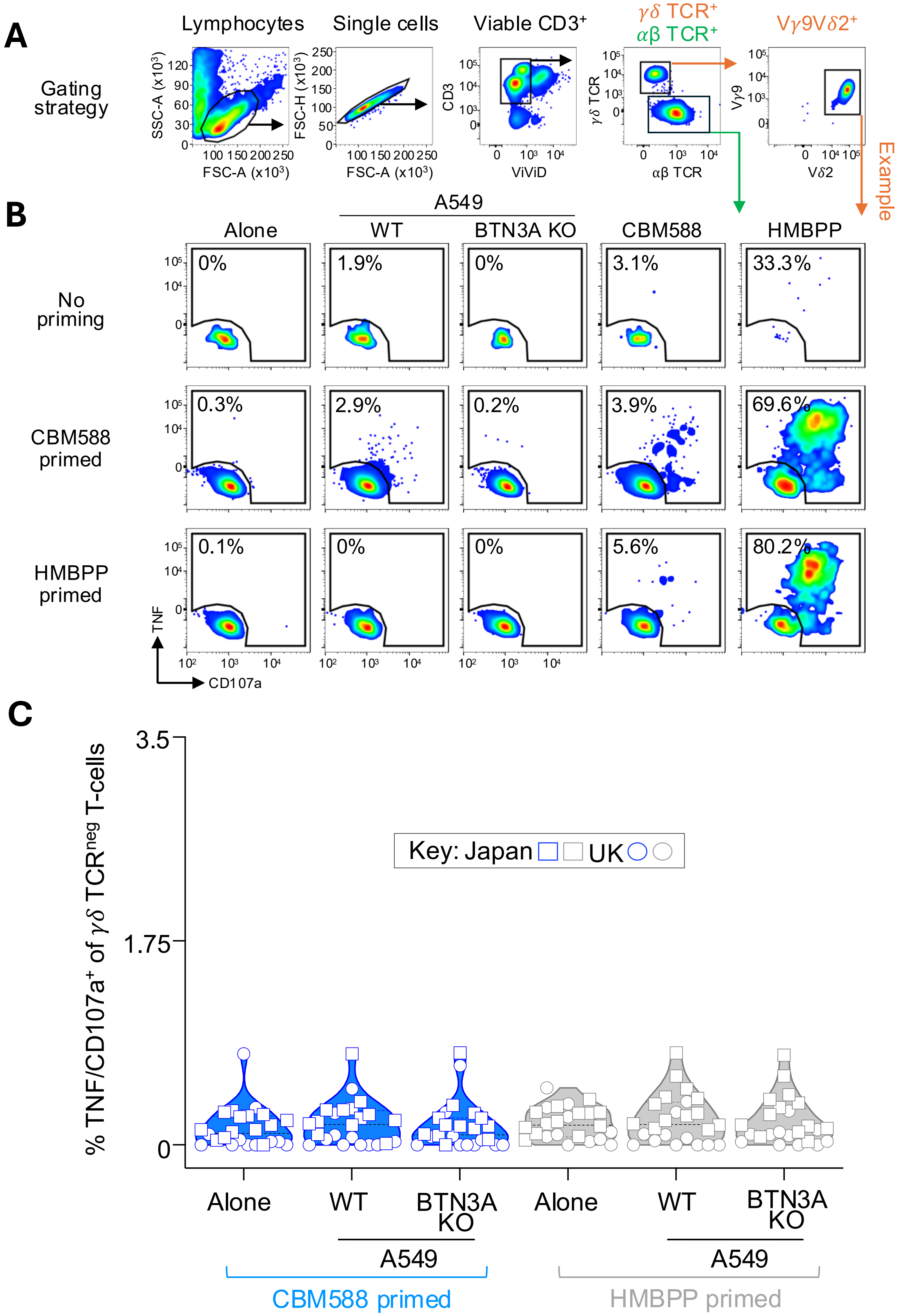


**Supplemental Figure 3. Gating strategy for functional testing of CBM588 tablet primed PBMCs. A**. T107 assay flow cytometry gating strategy for functional testing of CBM588 tablet (10 mg/mL) or HMBPP (10 ng/mL) primed PBMCs from healthy donors. **B**. Data set from a T107 assay for one healthy donor primed with CBM588 tablet or HMBPP, then tested against A549 cells, A549 BTN3A knockout cells, CBM588 tablet or HMBPP. Percentages shown for the TNF/CD107a$^+$ gate. **C**. $\alpha\beta$ TCR$^+$ T-cell reactivity (TNF/CD107a$^+$) of CBM588 tablet (10 mg/mL) or HMBPP (10 ng/mL) primed PBMCs from healthy donors from the United Kingdom or Japan (indicated in key) towards A549 and A549 BTN3A knock out (KO) cells.

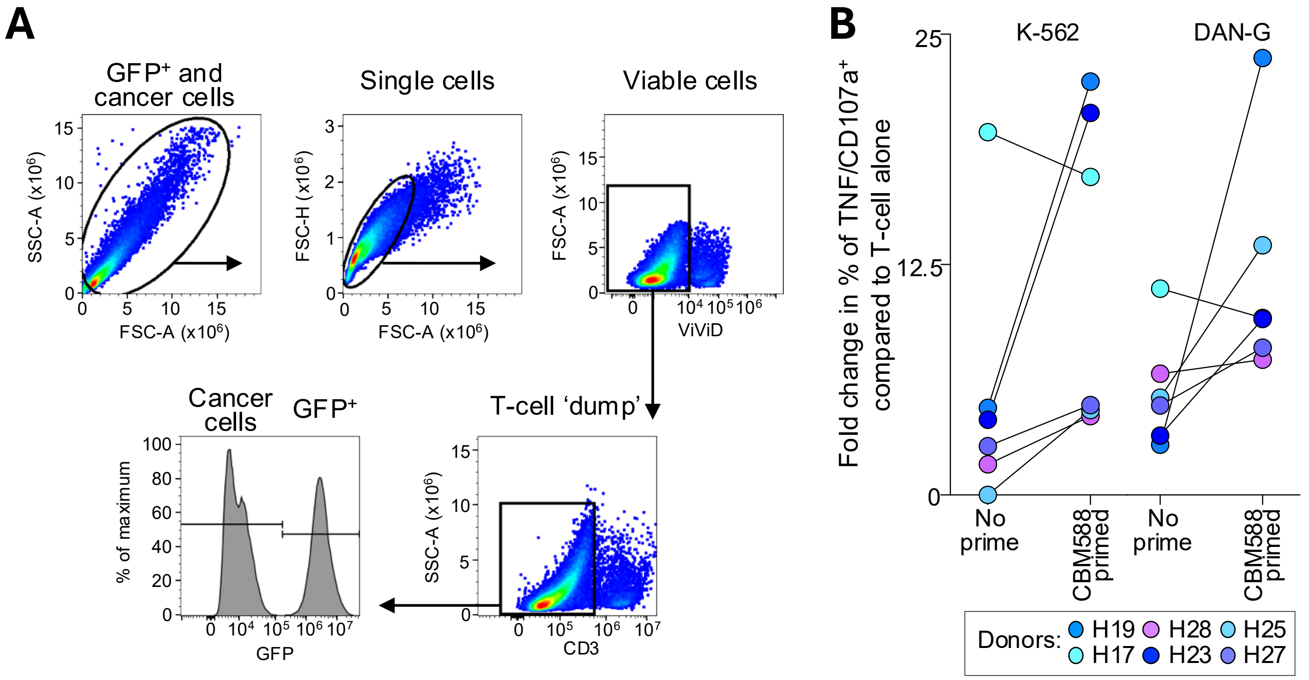


**Supplementary Figure 4. Gating strategy for flow cytometry based killing assays and data showing the recognition of cancer cells by CBM588 primed PBMCs. a**, Flow cytometry gating strategy for killing assays. GFP+ reference cells added to assay wells immediately prior to harvest and staining for flow cytometry. **b**, T107 assay (TNF and CD107a) for six healthy donors primed with CBM588 tablet then tested against K-562 (leukemia) and DAN-G (pancreatic) cancer cells. Gated on V$\gamma$9V$\delta$2+ T-cells. The fold increase in reactivity to cancer cells relative to T-cells alone is displayed.

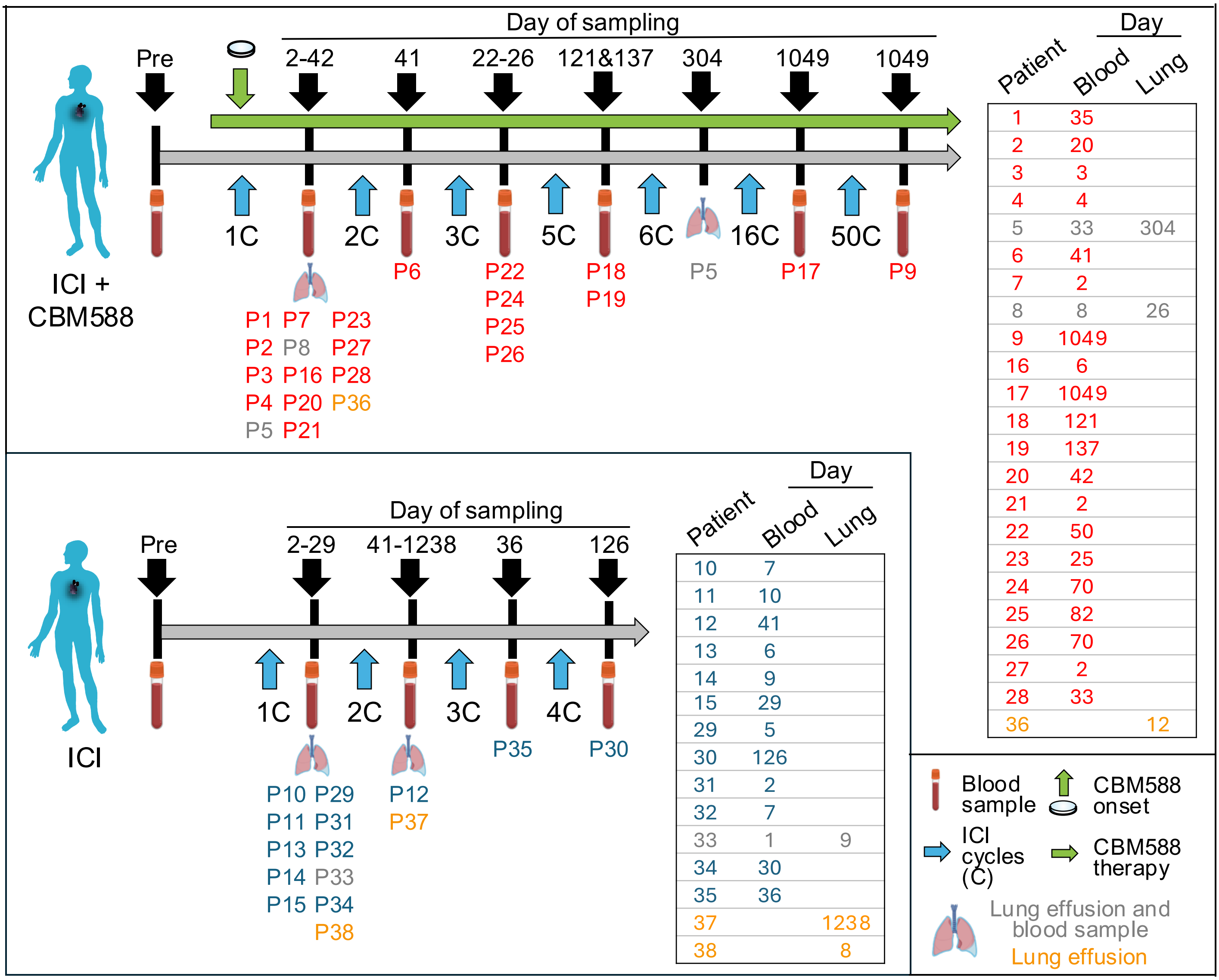


**Supplementary Figure 5. Patient overview.** Patients treated with immune checkpoint inhibitors (ICI), with or without CBM588. Blood (pre and/or post ICI) and lung pleural effusions taken post ICI (days), as indicated by the time courses and tables. ICI regimen: PD-1, + CTLA-4 (P15), or PD-L1 (P1 and P5).

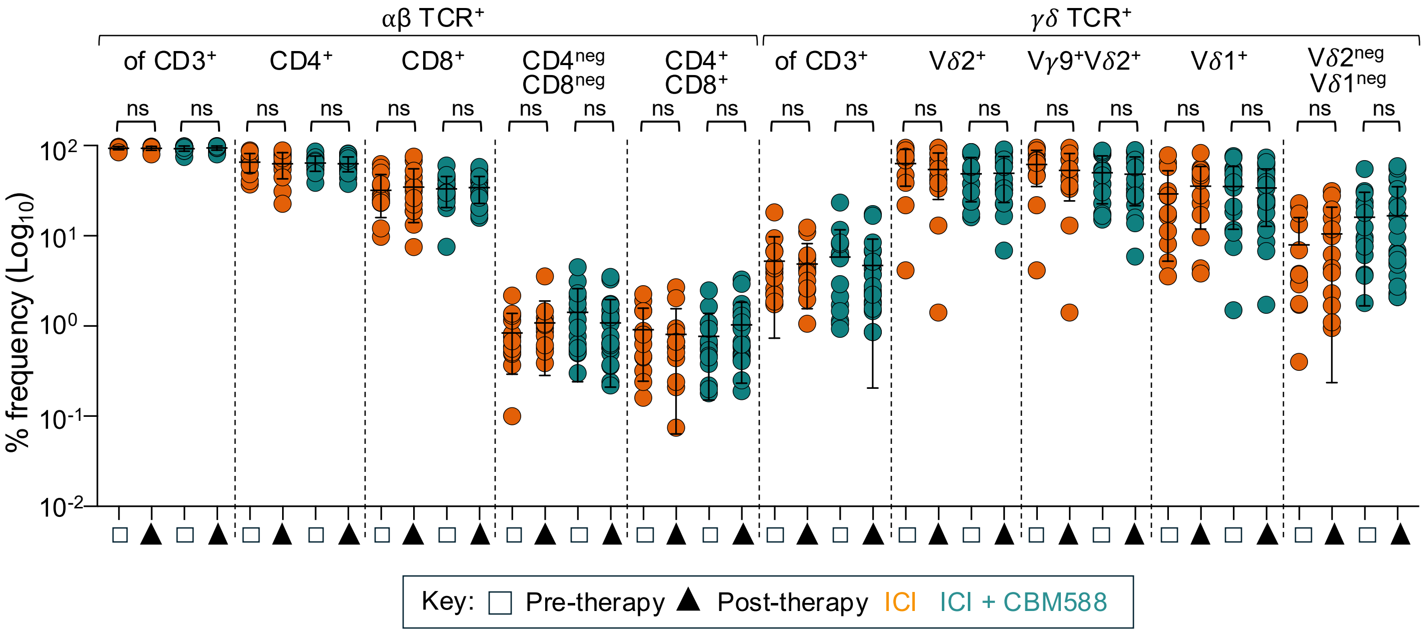


**Supplementary Figure 6. Frequency of peripheral blood T-cell subsets from lung cancer patients receiving ICI or ICI and CBM588.** Analysis of the frequency γδ TCR+ and αβ TCR+ T-cell subsets with statistical comparisons made between pre- and post-therapy for immune checkpoint inhibitor (ICI) therapy alone or ICI combined with CBM588.

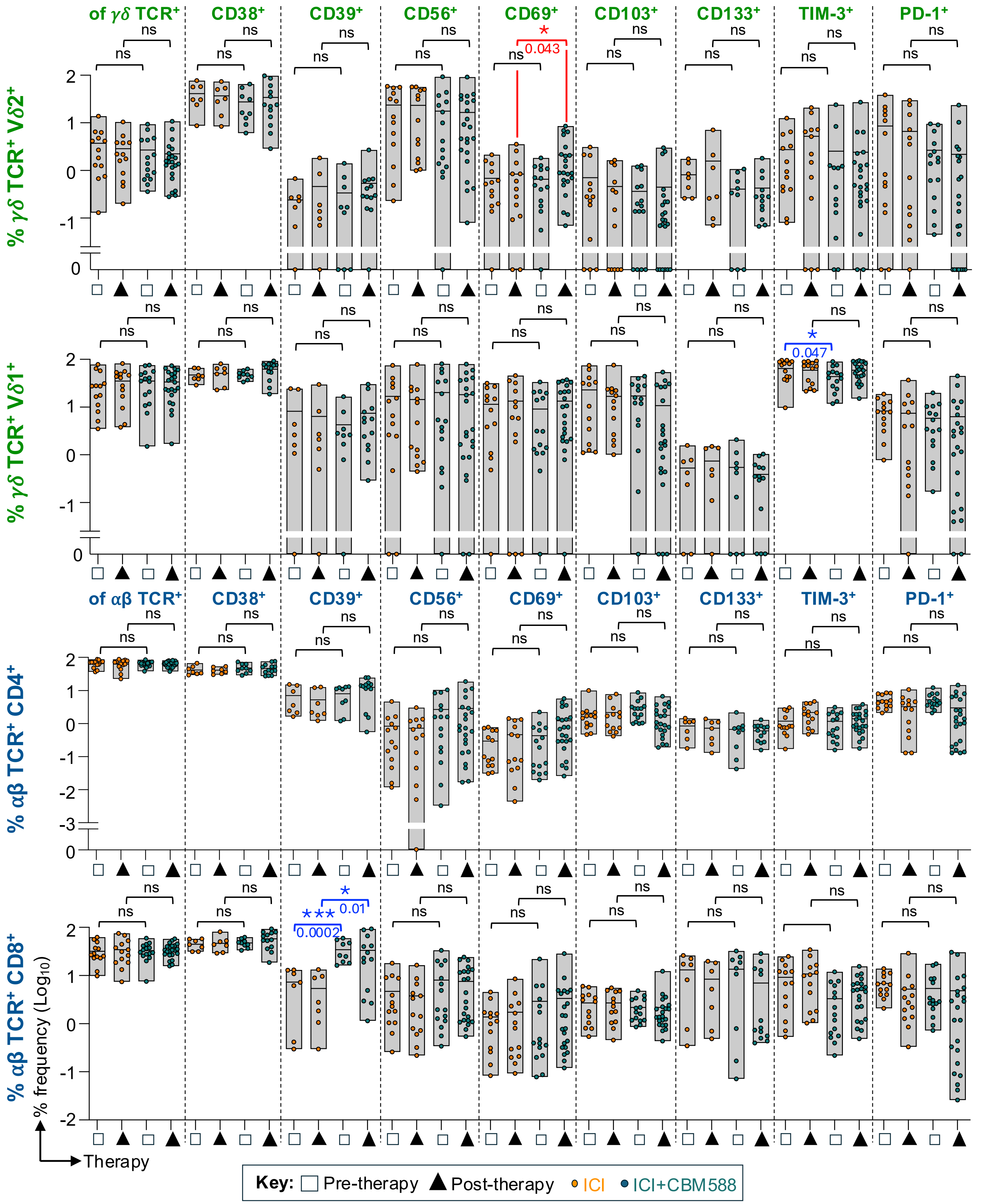


**Supplementary Figure 7. Frequency and phenotype of peripheral blood T-cell subsets from lung cancer patients receiving ICI or ICI and CBM588.** Analysis of the frequency and phenotype of various T-cell subsets, with statistical comparisons made between pre- and post-therapy for immune checkpoint inhibitor (ICI) therapy alone and ICI combined with CBM588. Notable findings include a significant difference in CD69 expression on Vδ2⁺ T-cells post-therapy between ICI and ICI+CBM588 (red text), whereas no significant difference was observed pre-therapy (same data as in **Fig. 3B**). For CD39 expression on αβ TCR⁺CD8⁺ T-cells, a significant difference for post-therapy CD39 between ICI and ICI+CBM588 was also seen before therapy (blue text). Additionally, TIM3 expression on Vδ1+ T-cells showed a significant difference only in the pre-therapy comparison between ICI and ICI+CBM588 (blue text). In summary, significant differences shown in blue text were present before therapy. The only significant difference present only post therapy is shown in red. P values are displayed for a two-tailed Mann-Whitney test.

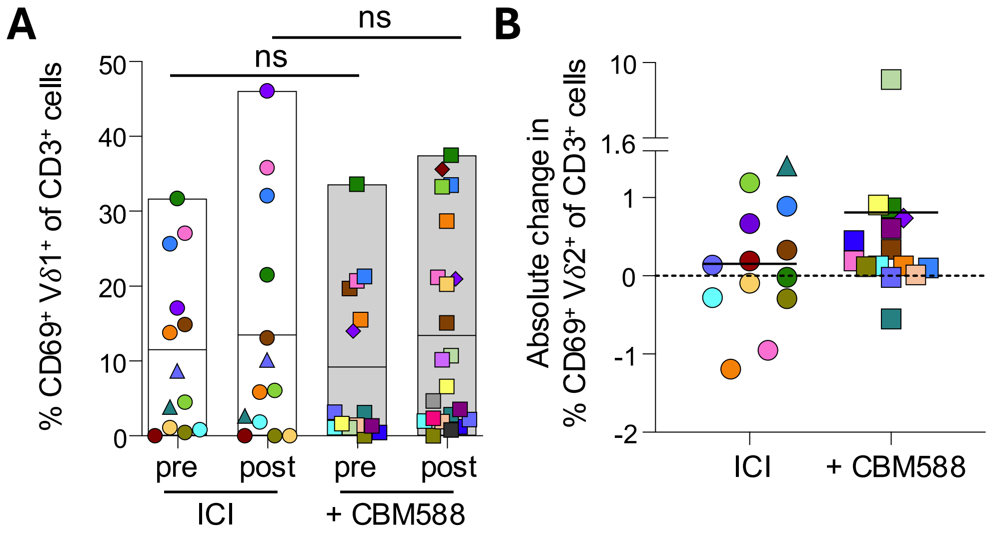


**Supplementary Figure 8. Pre- and post-therapy frequency of $CD69^+$ $V\delta1^+$ T-cells and post-therapy CD69 expression of peripheral blood $V\delta2^+$ T-cells. A**. Frequency of $CD69^+$ $V\delta1^+$ T-cells in $CD3^+$ T-cells from patients pre- and post-treatment with ICI, or ICI+CBM588. Statistical analysis by a Mann-Whitney two-tailed T-test. Same data as in **Supplementary Fig. 7**. Different symbol shapes and colors indicates individual patients (key in **Fig. 3A**). **B**. Absolute change (post- versus pre-therapy) in the frequency of peripheral blood $CD69^+$ $V\delta2^+$ T-cells for patients treated with ICI or ICI+CBM588. Solid line represents the mean and the dotted line is set at zero. Based on the data displayed in **Fig. 3C**. Different symbol shapes and colors indicates individual patients (key in **Fig. 3A**).

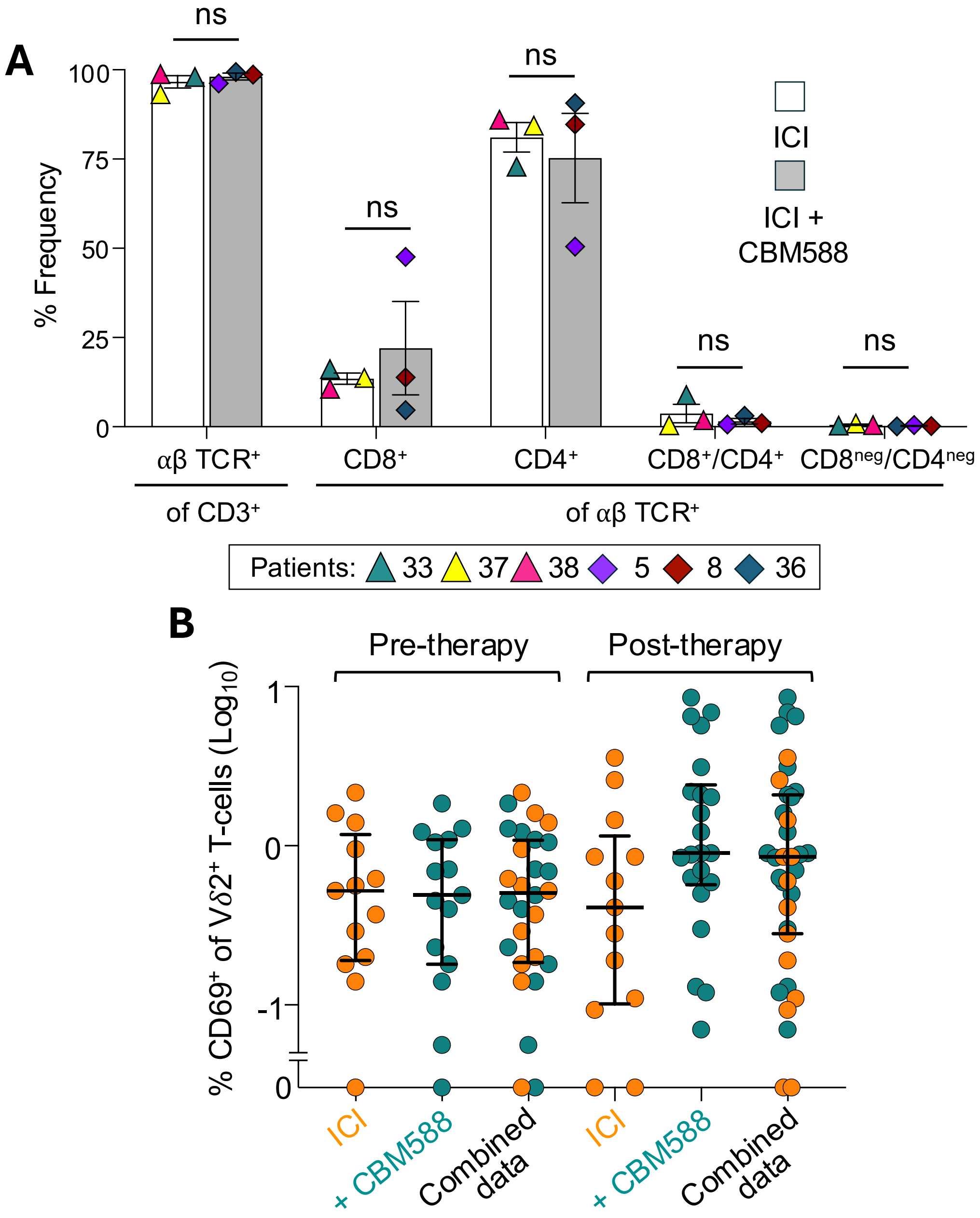


**Supplementary Figure 9. Frequency of T-cell subsets in pleural effusions and CD69 expression of peripheral blood V$\delta$2$^+$ T-cells from lung cancer patients treated with ICI or ICI+CBM588. A**. Percentage of αβ TCR$^+$ T-cell subsets in pleural effusions from lung cancer patients, treated with ICI or ICI+CBM588. Error bars depict standard error of mean. No significant difference was observed by paired two-tailed Student's t-test. **B**. CD69 expression on V$\delta$2$^+$ T-cells from the blood of patients on ICI or ICI+CBM588, pre- and post-therapy. Median is displayed with error bars depicting the interquartile range. The same data is also displayed in **Fig. 3B** and shown here for combined display of the cohorts with the median.

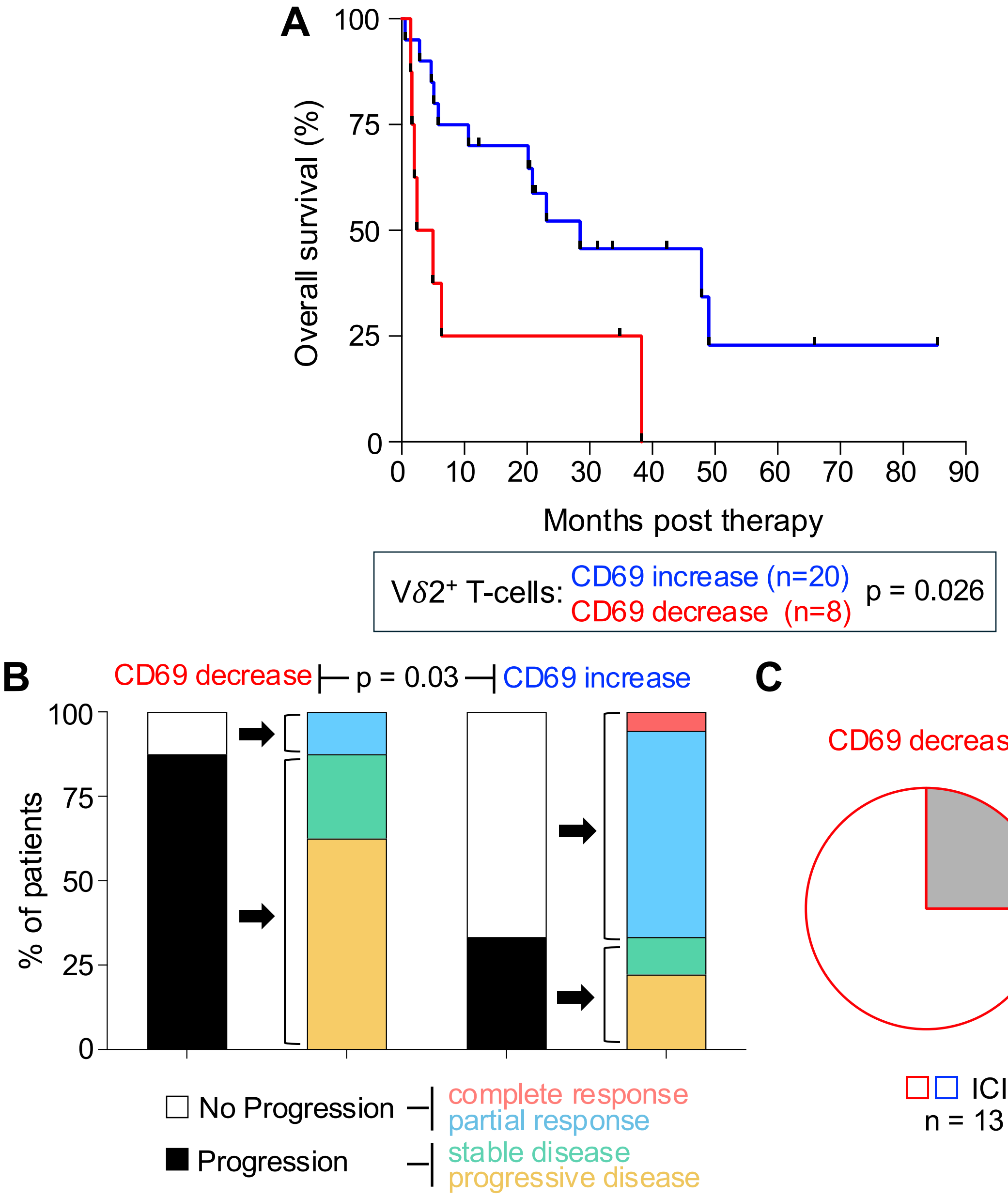


**Supplementary Figure 10. An increase in CD69 expression on V$\delta$2$^+$ T-cells post-therapy correlates with improved outcomes in lung cancer patients receiving immunotherapy. A**. Kaplan-Meier curve showing overall survival of lung cancer patients receiving either ICI or ICI+CBM588 (considered as one cohort, n = 28). Patients were categorized into two groups based on CD69 expression on V$\delta$2$^+$ T-cells: increase or decrease in CD69 post-therapy relative to pre-therapy. P value for a log-rank test. **B**. Decrease or increase in CD69 expression on V$\delta$2$^+$ T-cells categorized by clinical outcome according to the key. *p* value for Fisher's exact test. **C**. Patients on ICI or ICI + CBM588 within the CD69 decrease or increase categories.
